# Peroxisomal compartmentalization of amino acid biosynthesis reactions imposes an upper limit on compartment size

**DOI:** 10.1101/2023.03.06.531353

**Authors:** Ying Gu, Sara Alam, Snezhana Oliferenko

## Abstract

Cellular metabolism relies on just a few redox cofactors. Selective compartmentalization may prevent competition between metabolic reactions requiring the same cofactor. Is such compartmentalization necessary for optimal cell function? Is there an optimal compartment size? Here we probe these fundamental questions using peroxisomal compartmentalization of the last steps of lysine and histidine biosynthesis in the fission yeast *Schizosaccharomyces japonicus*. We show that compartmentalization of these NAD^+^ dependent reactions together with a dedicated NADH/NAD^+^ recycling enzyme supports optimal growth when an increased demand for anabolic reactions taxes cellular redox balance. In turn, compartmentalization constrains the size of individual organelles, with larger peroxisomes accumulating all the required enzymes but unable to support both biosynthetic reactions at the same time. We propose that compartmentalized biosynthetic reactions are sensitive to the size of the compartment, likely due to scaling-dependent changes within the system, such as enzyme packing density.

## Introduction

In eukaryotes, many metabolic pathways are partially or fully compartmentalized in subcellular organelles. Compartmentalization of enzymes and their reactants optimizes reaction flux within multiple metabolic networks and manages toxic effects of reaction by-products (Chen and Silver, 2012). The single membrane-bound peroxisomes exhibit great functional versatility. Despite peroxisomes sharing common biogenesis mechanisms and protein import machinery (Jansen et al., 2021), their enzymatic repertoire is highly variable among species, individual cell types and under different physiological conditions. For instance, in addition to well-established oxidative metabolism functions, such as fatty acid (FA) β-oxidation and hydrogen peroxide removal, the budding yeast peroxisomes participate in the biosynthesis of the amino acid L-lysine (Al-Saryi et al., 2017). Peroxisome-like organelles, glycosomes, in the Trypanosoma parasites, house enzymes for glycolysis, glycerol metabolism, pyrimidine and purine metabolism and ether-linked lipid synthesis (Hammond et al., 1985; Michels et al., 2006; Opperdoes and Borst, 1977; Pineda et al., 2018). During organismal development, peroxisomes may alter their functions by compartmentalizing distinct sets of enzymes. For instance, specialized peroxisomes, glyoxysomes, participate in converting storage lipids to carbohydrates in Arabidopsis seedlings, whereas in mature plants, peroxisomes harbouring a different repertoire of enzymes collaborate with other organelles in supporting photorespiration (Goto-Yamada et al., 2015; Oikawa et al., 2019).

Peroxisomes are a distributed organelle network, typically present as many individual compartments. Peroxisomes can arise *de novo* through the endoplasmic reticulum (ER) derived pre-peroxisomal vesicles (Agrawal and Subramani, 2016). Alternatively, they originate through fission of pre-existing mature peroxisomes, which is mediated by Pex11 that remodels the peroxisomal membrane and promotes activation of the dynamin-dependent membrane scission (Jourdain et al., 2008; Kuravi et al., 2006; Li and Gould, 2003; Williams et al., 2015). As peroxisomes grow and mature, cargo import into the peroxisomal matrix takes place. Delivery of proteins across the peroxisomal membrane is mediated by two cargo receptors binding to the specialized cis-motifs on cargo proteins – the PTS1-type receptor Pex5 and the PTS2-type receptor Pex7 (Gatto et al., 2000; Pan et al., 2013). Changes in peroxisome size and number have been associated with environmental stimuli in many organisms (Deb et al., 2022; Farre et al., 2022; Reddy et al., 1980; Reddy and Krishnakantha, 1975; Veenhuis et al., 1979; Veenhuis et al., 1987) and human disease (Jean Beltran et al., 2018; Legakis et al., 2002; Waterham et al., 2016).

The fission yeast *Schizosaccharomyces japonicus* provides an attractive stripped-down model for metabolic compartmentalization in peroxisomes. The fission yeast clade has lost the peroxisomal FA β-oxidation system. In fact, all fission yeasts lack a functional glyoxylate cycle, required for the conversion of acetyl-CoA generated by FA β-oxidation into four-carbon biosynthetic precursors, and so cannot use FAs as a carbon source (Rhind et al., 2011; Rutherford et al., 2022). Of note, the Lys3 and His2 enzymes catalysing the last steps of lysine and histidine biosynthesis, respectively, carry the C-terminal PTS1-like sequences (Rhind et al., 2011; Rutherford et al., 2022), suggesting that these metabolic reactions may take place in peroxisomes. *S. japonicus* grows at the cell tips and divides in the middle, maintaining a polarized pill-shape morphology throughout its vegetative growth (Gu and Oliferenko, 2019). *S. japonicus* maintains a large cell size and divides fast in the nutrient-rich medium but scales down its cell volume when switched to the minimal medium, where cells are forced to synthesize all amino acids (Gu and Oliferenko, 2019). Does peroxisomal compartmentalization of amino acid biosynthesis facilitate metabolic adaptation in nutrient-poor conditions? Given that the efficiency of compartmentalized reactions may depend on the size of the compartment (Hinzpeter et al., 2017), are there specific size requirements for optimal biosynthetic output? Here we probe these fundamental questions using *S. japonicus* as a model system.

## Results

### The glycerol-3-phosphate dehydrogenase Gpd2 provides NAD^+^ for histidine and lysine biosynthesis in peroxisomes

*S. japonicus* does not respire oxygen, unlike *S. pombe*, but grows fast and maintains high energy content. Rather than using the electron transport chain, it relies on the reduction of dihydroxyacetone phosphate (DHAP) to glycerol-3-phosphate (G3P) by the cytosolic glycerol-3-phosphate dehydrogenase Gpd1 to oxidize NADH, to sustain purely fermentative growth (Alam et al., 2022). *S. japonicus* but not *S. pombe* cells lacking Gpd1 exhibit severe growth defects and cell shape abnormalities in the rich YES medium and are unable to grow at all in the minimal EMM medium (Fig. 1A, C, Fig. S1A and (Alam et al., 2022)). The fission yeast genomes encode an additional NADH-dependent glycerol-3-phosphate dehydrogenase, Gpd2 (Rutherford et al., 2022; Yamada et al., 1996). *S. japonicus gpd2*Δ cells exhibited normal morphology, but their rate of growth was slightly attenuated in the rich medium (Fig. 1B-D). Interestingly, the mutant cells rounded up and fully arrested growth upon transfer to EMM (Fig. 1B-D). The loss of cellular polarity leading to virtually spherical cells is typical for growth arrest in *S. japonicus* that relies on its geometry for cell division (Gu and Oliferenko, 2019) and can be used as a proxy for estimating population growth with a single-cell resolution. The inability of Gpd2-deficient cells to grow in the minimal medium was conserved in *S. pombe*, consistent with a previous report (Fig. S1A-B and (Yamada et al., 1996)).

Gpd2 was enriched in peroxisomes in *S. japonicus* (Fig. 1E, left panel). As expected from the presence of the C-terminal PTS1-type peroxisome targeting signals, Lys3 and His2 enzymes catalysing the last reactions of lysine and histidine biosynthesis respectively, also localised to peroxisomes (Fig. 1E, middle and right panels). Both reactions require NAD^+^, suggesting a possible role for Gpd2 in re-oxidizing NADH to sustain lysine and histidine synthesis in peroxisomes (Fig. 1F). Indeed, growth inhibition of *S. japonicus gpd2*Δ cells was rescued only when the minimal medium was supplemented with both lysine and histidine (Fig.1D).

Growth in minimal medium requires cells to synthesize all amino acids. Since many anabolic reactions require NAD^+^, it may challenge the cellular redox balance. Indeed, the NAD^+^/NADH ratio dramatically decreased when *S. japonicus* cells grown in the rich YES medium were transferred to the minimal EMM (Fig. 1G). Consistent with a shift in cellular redox status, the switch to the minimal medium elicited transient upregulation of cytosolic catalase Ctt1, a hallmark of oxidative stress response (Nakagawa et al., 1995) (Fig. S1C, D). Such a dramatic upregulation did not occur in *S. pombe* (Fig. S1D, upper panel). This result indicates that redox balancing mechanisms in *S. japonicus* may struggle to meet the demands of increased anabolism.

The catalase upregulation was even more pronounced in *S. japonicus* cells lacking Gpd2 (Fig. 1H and Fig. S1E, upper panel). Supplementing the growth medium with lysine and histidine restored the kinetics of catalase regulation in *gpd2*Δ cells to wild type levels (Fig. 1H and Fig. S1E, bottom panel). Gpd1 also exhibited major upregulation in *S. japonicus* upon the shift to the minimal medium, in line with its function in redox balancing (Fig. S1D, lower panel). In fact, an ectopic copy of Gpd1 fused to PTS1, Gpd1^PTS1^, and expressed from the *gpd2* promoter, was able to rescue polarity scaling and growth in *S. japonicus gpd2*Δ cells upon shift to the minimal medium (Fig. 1I, J). Taken together, these results suggest that beyond the role in synthesizing glycerol-3-phosphate, the glycerol-3-phosphate dehydrogenases Gpd1 and Gpd2 execute redox balancing in the cytosol and inside peroxisomes, respectively.

**Figure 1.**
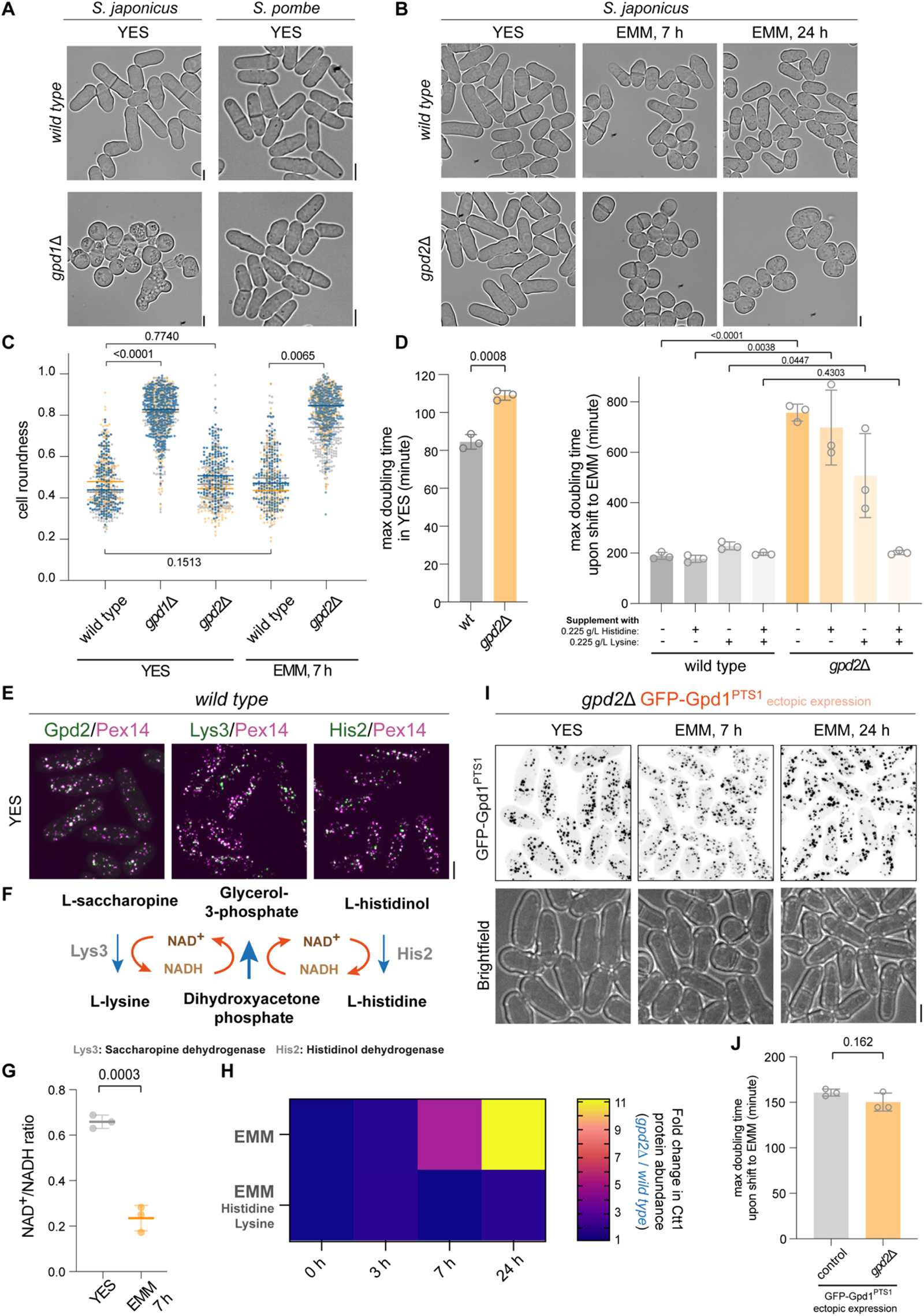
Peroxisomal glycerol-3-phosphate dehydrogenase Gpd2 enables *de novo* lysine and histidine biosynthesis. **(A)** Brightfield images of wild type and *gpd1*Δ *S. japonicus* and *S. pombe* cells grown in rich YES medium. (**B**) Brightfield images of wild type and *gpd2*Δ *S. japonicus* cells in YES (left panel) and following a switch to the minimal EMM medium for 7 and 24 hours. (**C**) Cell morphology profiles (using a ‘cell roundness’ metric) of wild type, *gpd1*Δ and *gpd2*Δ *S. japonicus* cultures in indicated media. Bars represent population medians. (**D**) Growth rates of wild type and *gpd2*Δ *S. japonicus* cultures grown in YES (left) and post medium switch to EMM, with or without supplementation with indicated amino acids (right). (**E**) Colour overlays of maximum Z-projection spinning disk confocal images of *S. japonicus* cells co-expressing Pex14-mCherry (magenta) and Gpd2-mNeonGreen, GFP-Lys3 or GFP-His2, respectively (green). (**F**) Illustration of the last steps in L-lysine and L-histidine biosynthesis pathways, catalysed by Lys3 and His2 enzymes, respectively. Both reactions reduce the cofactor NAD^+^ to NADH, which can be re-oxidised to NAD^+^ via glycerol-3-phosphate (G3P) synthesis from dihydroxyacetone (DHAP). (**G**) Total cellular NAD^+^/NADH ratios of *S. japonicus* grown in YES and 7 hours post-switch to EMM. (**H**) Catalase Ctt1-mNeonGreen protein abundance (measured as average intensity of whole cell Z-projections) in *gpd2*Δ *S. japonicus* cells normalised to the wild type, at 0, 3, 7 and 24 hour time points, post-switch from YES to EMM (top) or EMM supplemented with L-lysine and L-histidine (bottom). Heatmap shows ratios between populational means of average cell fluorescence intensities. (**I**) Maximum Z-projection spinning disk confocal images of *gpd2*Δ *S. japonicus* cells expressing an extra copy of GFP-tagged Gpd1 with the PTS1 targeting signal, in indicated media. Brightfield images are shown underneath the corresponding fluorescence channel images. (**J**) Growth rates of *S. japonicus* wild type and *gpd2*Δ cells expressing an extra copy of GFP-tagged Gpd1-PTS1, post-switch to EMM. (**A**, **B, E, I**) Scale bars represent 5 μm. (**C, D, G, J**) *p* values are derived from unpaired t-test. (**C**, **D, G, H**, **J**) Values are derived from three biological replicates. (**D, G, J**) Bars represent mean values ±SD.

**Supplemental Figure 1.**
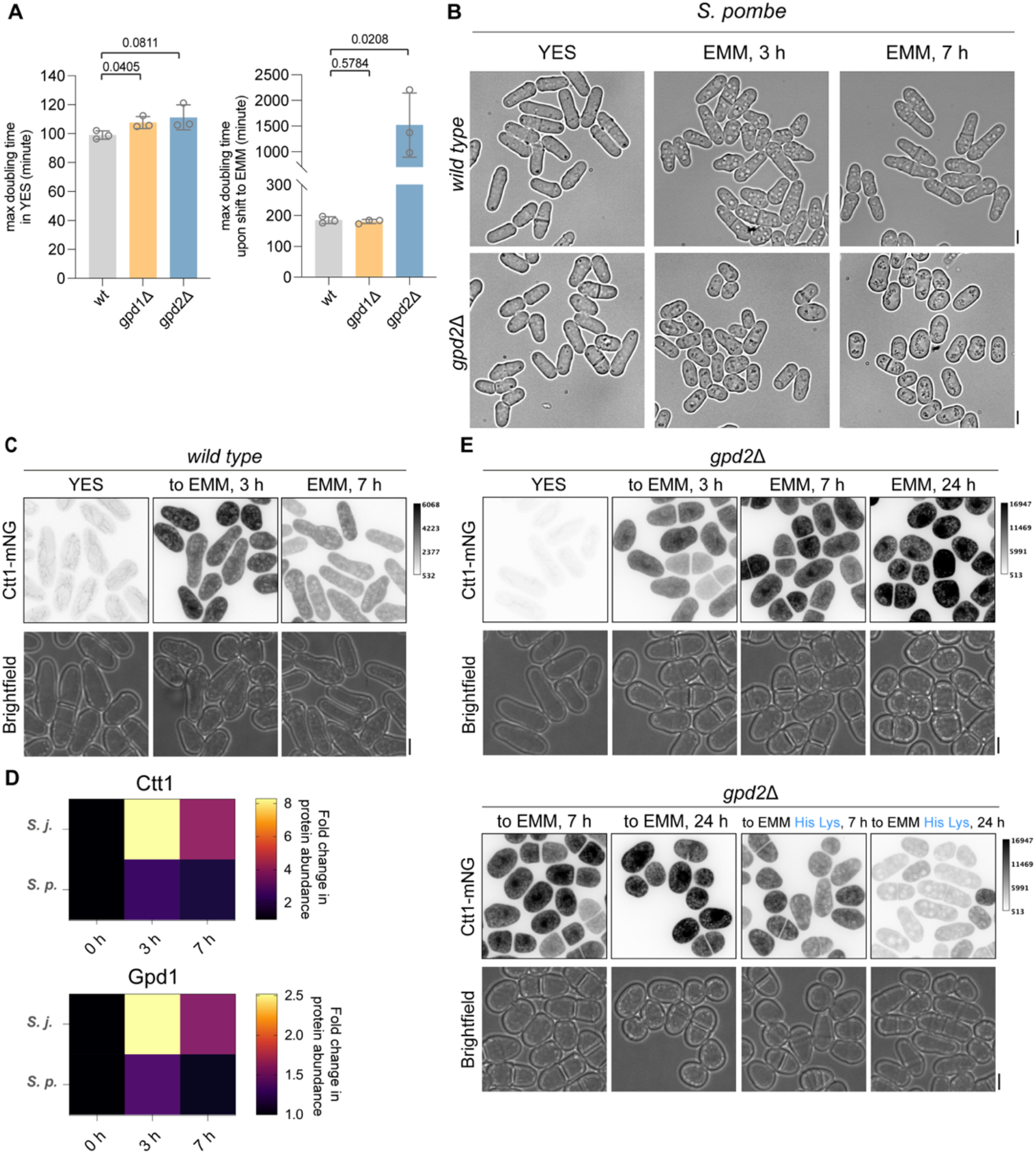
**(A)** Growth rates of wild type, *gpd1*Δ and *gpd2*Δ *S. pombe* cultures grown in YES (left) and post-shift to EMM (right). (**B**) Brightfield images of *w*ild type and *gpd2*Δ *S. pombe* cells in YES (left panel) and following a switch to the minimal EMM medium for 3 hours and 7 hours. *p* values are derived from unpaired t-test analysis. Bars represent mean values ±SD. (**C**) Maximum Z-projection spinning disk confocal images of *S. japonicus* expressing Ctt1-mNeonGreen in indicated conditions. (**D**) Catalase Ctt1-mNeonGreen (*top*) and Gpd1 (*bottom*) protein abundance (measured as average intensity of whole cell Z-projection) in *S. japonicus* and *S. pombe* wild type cells sampled at 0, 3 and 7 hour time points post-shift to EMM. Heatmap shows ratios between populational means of average cell fluorescence intensity in each dataset. (**E**) Maximum Z-projection spinning disk confocal images of *gpd2*Δ *S. japonicus* cells expressing Ctt1-mNeonGreen in indicated conditions. Supplemented amino acids L-lysine (Lys) and L-histidine (His) are indicated in blue. Brightfield images are shown underneath the corresponding fluorescence channel images. (**B, C, E**) Scale bars represent 5 μm. (**A**, **D**) Values are derived from three biological replicates. (**C**, **E**) Greyscale calibration bars are shown, note much higher fluorescence levels of Ctt1-mNeonGreen in (**E**).

### Selective compartmentalization of Lys3 and His2 facilitates cellular redox balance to meet the demand for *de novo* amino acid biosynthesis

To understand why cells may benefit from selective compartmentalization of certain amino acid biosynthesis reactions, we sought to create scenarios, where NAD^+^ dependent Lys3 and His2 enzymes normally present in peroxisomes were delocalized to the cytosol in the presence or absence of the glycerol-3-phosphate dehydrogenases.

*S. japonicus* cells lacking Pex5 but not Pex7 failed to import Lys3 and His2, both of which have C-terminal PTS1, into peroxisomes. Similarly, the ectopically expressed Gpd1^PTS1^ delocalized to the cytosol in *pex5*Δ cells (Fig. S2A). The *S. japonicus* Gpd2 harbours an N-terminal PTS2; as expected, it largely redistributed to the cytoplasm in the absence of Pex7 (Fig. S2A). Either the deletion or point mutations of PTS2 (see Experimental Procedures) fully delocalized Gpd2 to the cytoplasm and arrested growth of mutant cells upon transfer to the minimal medium (Fig. S2B), underscoring functional importance of peroxisomal targeting of this protein. Interestingly, Gpd2 import to peroxisomes was also partially disrupted by Pex5 loss of function (Fig. S2A), consistent with previous reports in mammalian cells, plants, *Trypanosoma* and the fungal plant pathogen *Ustilago maydis*, where Pex5 was shown to serve as a Pex7 co-receptor for the PTS2 cargo translocation across the peroxisomal membrane (Ast et al., 2022; Galland et al., 2007; Hayashi et al., 2005; Otera et al., 1998; Woodward and Bartel, 2005).

We wondered if releasing Lys3 and His2 from peroxisomes into cytosol by disabling Pex5 would allow lysine and histidine biosynthesis utilizing cytosolic NAD^+^. Indeed, not only *pex5*Δ but also *pex5*Δ *gpd2*Δ double mutant *S. japonicus* cells were capable of growth in the minimal medium, albeit at a reduced growth rate (Fig. 2A, C and Fig. S2C). We also observed a similar rescue of growth without amino acid supplementation when we deleted *pex3*, which fully abolishes peroxisome biogenesis, or *pex3* in combination with *gpd2* (Fig. S2C). Interestingly, *pex3* deletion also noticeably improved the growth of *gpd1*Δ cells in the rich YES medium but not in the minimal medium where cells are forced to synthesize all amino acids (Fig. 2B; see Fig. 1C for comparison). This result argues that although in principle, Gpd2 can substitute for Gpd1 in supporting redox balance in the cytosol, the correct dosage of NAD^+^ regenerating enzymes is critical when cellular anabolic demands are increased.

Gas chromatography-mass spectrometry (GC-MS) measurements showed significant disbalance in the intracellular levels of many amino acids in *pex5*Δ *S. japonicus* cells grown in the minimal medium, as compared to the wild type (Fig. 2D). The levels of many amino acids synthesized in the cytosol were decreased (Fig. 2D, E). In particular, this was the case for the amino acids the synthesis of which requires NAD^+^ or NADPH. Leucine but not valine levels were decreased in *pex5*Δ cells, as compared to the control (Fig. 2D, E). Both amino acids share the same precursor, pyruvate, and their production pathways are linked within mitochondria (Ohtsuka et al., 2022). Yet, only the leucine biosynthesis pathway utilises a cytosolic NAD^+^-dependent enzyme, Leu1, in contrast to the valine production, which requires the NADPH cofactor for the Ilv5-mediated reaction in mitochodria (Kikuchi et al., 1988; Ohtsuka et al., 2022; Petersen and Holmberg, 1986; Rhind et al., 2011; Rutherford et al., 2022).

Strikingly, the intracellular levels of lysine were profoundly increased in *pex5*Δ cells (Fig. 2D, E). In budding yeast, the lack of the peroxisomal E3 ubiquitin ligase Pex12, which delocalizes the PTS1 cargo to the cytosol, leads to transcriptional upregulation of several genes in the lysine biosynthesis pathway (Fig. S2D) (Breitling et al., 2002). The upregulation was proposed to be a compensation response regulated by the transcription activator Lys14. However, Lys14 is not conserved outside of Saccharomycetaceae, and we did not observe dramatic changes in Lys2, Lys4, Lys9 and Lys12 protein levels in *pex5*Δ *S. japonicus* mutants (Fig. S2E, F). Of note, the *S. japonicus* Lys4 was confined to mitochondria, unlike its orthologs in *S. pombe* (Fig. S2G) or *S. cerevisiae* (Breker et al., 2014; Breker et al., 2013; Chen et al., 1997; Feller et al., 1999). Our results suggest that upregulation of lysine production by delocalized Lys3 could drive higher flux through the entire lysine biosynthesis pathway, which includes several NAD^+^ and NADPH-dependent reactions, potentially competing with other metabolic reactions and causing amino acid disbalance (Fig. 2D, E).

Since growth in the minimal medium taxes the intracellular supply of NAD^+^ (Fig. 1G), we wondered if providing cells with more NADH-oxidizing activity could rescue poor growth of *pex5*Δ cells in EMM. We made use of an ectopically expressed PTS1-carrying version of Gpd1 (Gpd1^PTS1^) that localized to the cytosol upon loss of Pex5 cargo receptor (Fig. S2A). Indeed, *pex5*Δ cells expressing ectopic Gpd1^PTS1^ proliferated normally in the minimal medium (Fig. 2F). Although the amino acid profile of *pex5*Δ cells expressing ectopic Gpd1^PTS1^ was now more similar to the control, we did not observe a full rescue (Fig. 2G). In particular, lysine levels remained high, suggesting that the compartmentalization of the last step of lysine biosynthesis in peroxisomes was important for controlling the activity of this biosynthetic pathway.

**Figure 2.**
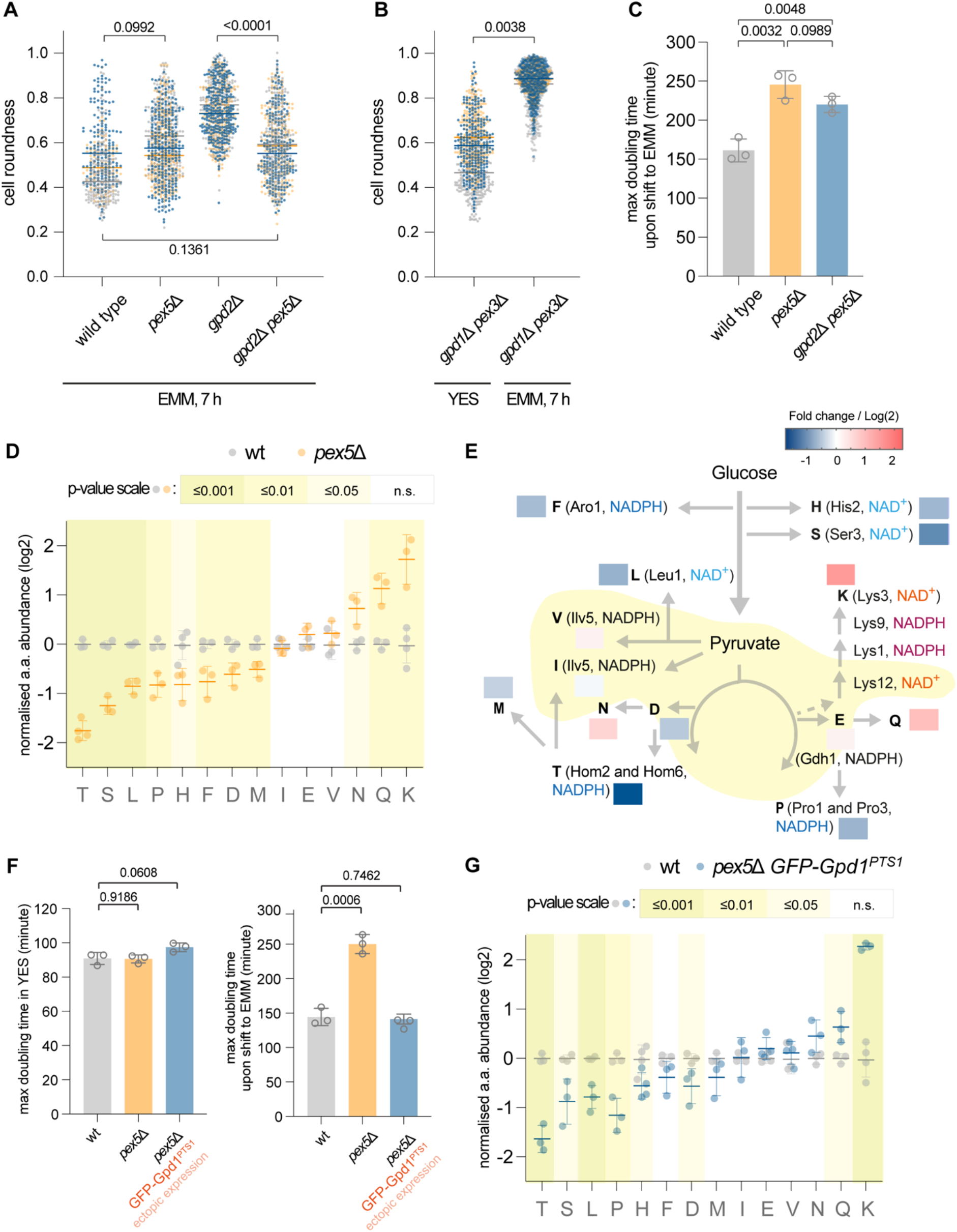
Compartmentalization of Lys3 and His2 enzymes in peroxisomes satisfies the redox demand for balanced amino acid biosynthesis. (**A**, **B**) Cell morphology profiles (using a ‘cell roundness’ metric) of *S. japonicus* cells of indicated genotypes under indicated conditions. (**C**) Growth rates of *S. japonicus* cultures of indicated genotypes post medium switch to EMM. (**D**) Fold changes of amino acids abundances (expressed as log2) in *pex5*Δ *S. japonicus* cells normalised to the wild type, sampled at 7 hours post-shift to EMM. (**E**) Illustration of amino acid biosynthesis pathways derived from glycolysis and the TCA cycle. Pathway logics are indicated by grey lines with arrows. Please note that the TCA pathway operates in a bifurcated configuration in *S. japonicus* (Alam et al., 2022). Reactions localized in mitochondria are shaded in yellow. Fold changes of individual amino acid abundance in *pex5*Δ *S. japonicus* cells normalised to the wild type shown in (**D**) are represented as heatmaps. Known enzymes involved in redox reactions utilising either NAD^+^ or NADPH co-factors are indicated next to respective amino acid products. (**F**) Growth rates of *S. japonicus* wild type, *pex5*Δ and *pex5*Δ cells expressing an extra copy of GFP-tagged Gpd1-PTS grown in YES (left) and post-shift to EMM (right). (**G**) Fold changes of amino acids abundances (expressed as log2) in *pex5*Δ *S. japonicus* cells expressing an extra copy of GFP-tagged Gpd1-PTS1 normalised to the wild type, sampled at 7 hours post-shift to EMM. (**A**-**C**, **F**) Values are derived from three biological replicates. *p* values are derived from unpaired t-test. (**D**, **G**) Values are from two technical repeats of two biological replicates. *p* values are derived from unpaired t-test analysis and colour-coded (see the scale above the graphs). (**A, B**) Bars represent mean population medians. (**C**, **D, F, G**) Bars represent mean values ±SD.

**Supplemental Figure 2.**
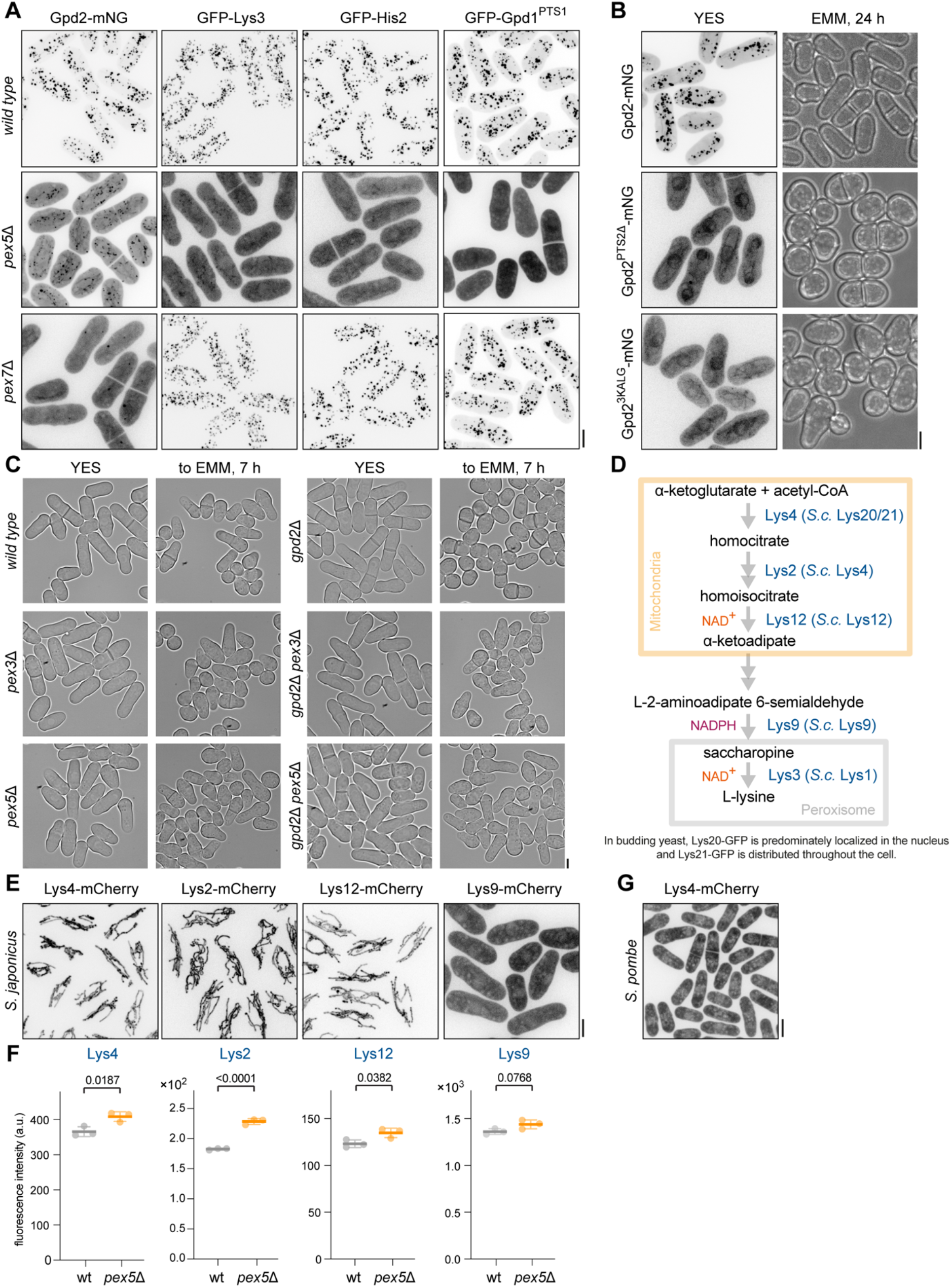
**(A)** Maximum Z-projection spinning disk confocal images of wild type, *pex5*Δ and *pex7*Δ *S. japonicus* cells grown in YES, expressing Gpd2-mNeonGreen, GFP-Lys3, GFP-His2 or GFP-Gpd1^PTS1^, respectively. (**B**) Maximum Z-projection spinning disk confocal images of *S. japonicus* cells expressing mNeonGreen-tagged Gpd2 constructs: wild type Gpd2, a short N-terminal truncation removing PTS2 (Gpd2^PTS2Δ^), and Gpd2 carrying point mutations in PTS2, which abolish peroxisomal import (Gpd2^3KALG^) under indicated conditions. (**C**) Brightfield images of *S. japonicus* cells of indicated genotypes grown in YES and post-shift to EMM for 7 hours. (**D**) Illustration of L-lysine biosynthesis pathway, catalysed by enzymes labelled in dark blue. Assignment of cellular compartmentalisation of these enzymes is based on fluorescent protein tagging shown in (**E**) and Fig. 1E. *S. cerevisiae* (*S. c*.) homologs are indicated. Mitochondria are indicated in yellow and peroxisomes in grey. (**E**) Maximum Z-projection spinning disk confocal images of *S. japonicus* cells expressing mCherry-tagged Lys4, Lys2, Lys12 and Lys9. (**F**) Protein abundance (measured as average intensity of whole cell Z-projection) of mCherry-tagged enzymes shown in (**E**) in *S. japonicus* wild type and *pex5*Δ cells sampled at 7 hours post-shift to EMM. Values are derived from three biological replicates. *p* values are derived from unpaired t-test analysis. Bars represent mean values ±SD. (**G**) Maximum Z-projection spinning disk confocal images of *S. pombe* cells expressing mCherry-tagged Lys4. (**A**-**C, E, G**) Scale bars represent 5 μm.

### Abnormal peroxisome compartment architecture impedes lysine and histidine biosynthesis

We noticed that the *S. japonicus* cells lacking the peroxisomal fission factor Pex11 (Li and Gould, 2002) struggled to adapt to growth in the minimal EMM medium (Fig. 3A, B, compare with Fig. 1B, C). Supplementation of the medium with either lysine or histidine significantly restored growth and normal cell morphology in *pex11*Δ cultures (Fig. 3B). Hyper-accumulation of the cytosolic catalase Ctt1, indicative of prolonged oxidative stress response was also alleviated in *pex11*Δ cells when the growth medium was supplemented with either lysine or histidine (Fig. 3C, D). These results suggested that although Lys3 or His2-mediated reactions could work in principle, they were not sufficiently productive when the demand for lysine and histidine biosynthesis was high. Interestingly, some *pex11*Δ cells were able to return to polarized growth after prolonged incubation in the EMM (see the 24-hour time-point in Fig. 3A). The ability to grow in the minimal medium coincided with a virtual disappearance of peroxisomes in these cells (Fig. S3A, B), likely due to a failure in the inheritance of abnormally large peroxisomes by some daughter cells.

Targeting of ectopic Gpd1^PTS1^ to peroxisomes was unable to rescue the growth arrest of *pex11*Δ cells (Fig. 3E), suggesting that the insufficient dosage of the redox balancing Gpd2 enzyme was not responsible for functional lysine and histidine auxotrophy in this genetic background.

We concluded that other parameters pertaining to peroxisome compartment architecture, such as the size and/or the number of individual peroxisomes may dictate the productivity of lysine and histidine biosynthesis.

**Figure 3.**
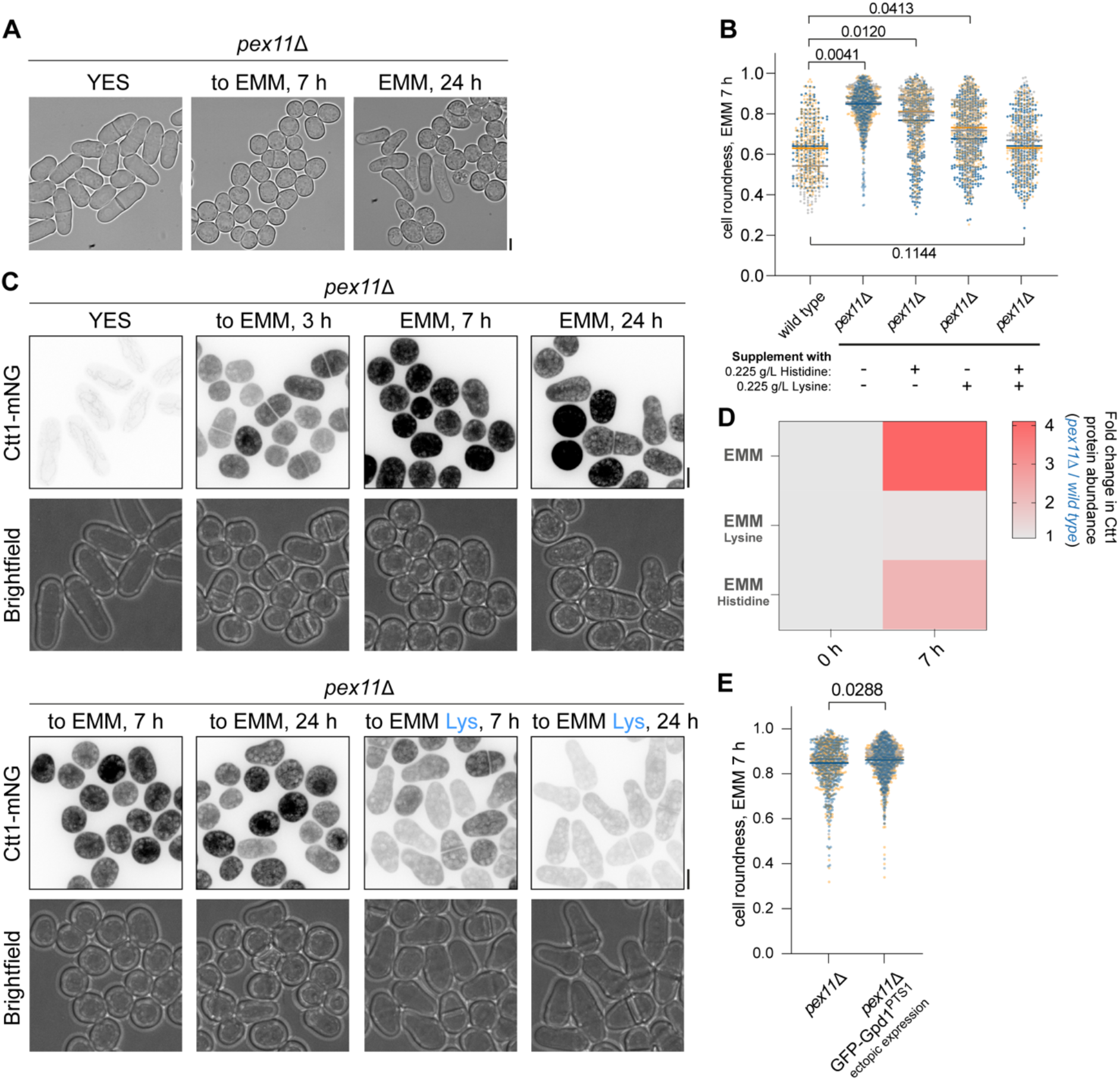
Abnormal peroxisome compartment architecture blocks lysine and histidine biosynthesis. **(A)** Brightfield images of *pex11*Δ *S. japonicus* cells grown in YES and post-shift to EMM. (**B**) Cell morphology profiles (using a ‘cell roundness’ metric) of wild type and *pex11*Δ *S. japonicus* cells sampled at 7 hours post-shift to EMM or EMM with indicated amino acid supplementation. **(B)** Maximum Z-projection spinning disk confocal images of *pex11*Δ *S. japonicus* cells expressing Ctt1-mNeonGreen in indicated growth conditions. Supplemented amino acids L-lysine (Lys) and L-histidine (His) are indicated in blue. Brightfield images are shown underneath the corresponding fluorescence channel images. (**D**) Catalase Ctt1-mNeonGreen protein abundance (measured as average intensity of whole cell Z-projection) in *pex11*Δ *S. japonicus* cells normalised to the wild type, at time points 0, 3, 7 and 24 hours, post-switch from YES to EMM (top) or EMM supplemented with L-lysine (bottom). Heatmap shows ratios between populational means of average cell fluorescence intensity. Values are derived from three biological replicates. (**E**) Cell morphology profiles (using a ‘cell roundness’ metric) of *pex11*Δ and *pex11*Δ cells expressing an extra copy of GFP-Gpd1^PTS1^, sampled at 7 hours post-shift to EMM. (**A, C**) Scale bars represent 5 μm. (**B**, **D**, **E**) Values are derived from three biological replicates. (**B**, **E**) Bars represent mean values ±SD. *p* values are derived from unpaired t-test analysis.

**Supplemental Figure 3.**
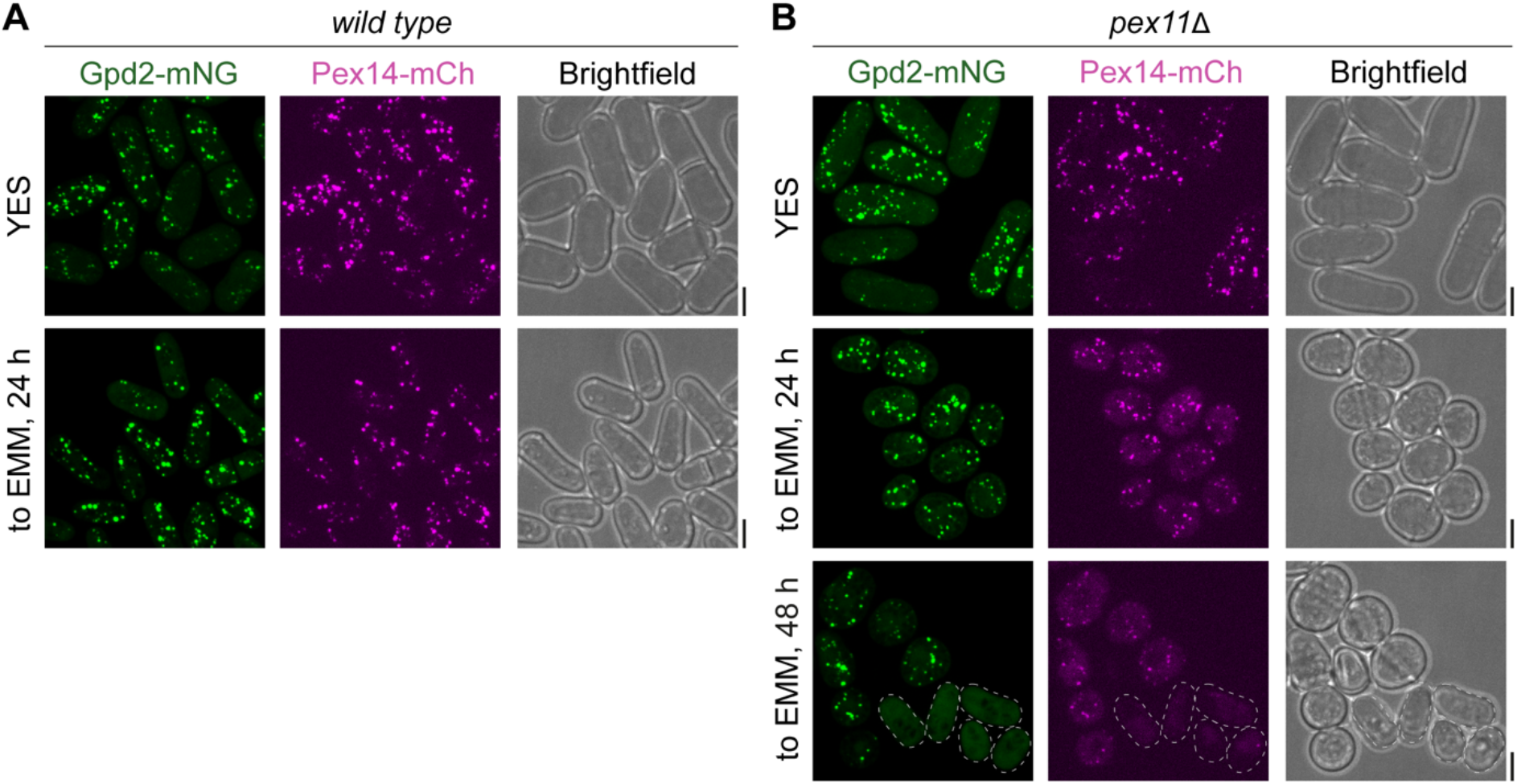
**(A)** Maximum Z-projection spinning disk confocal images of *S. japonicus* wild type cells co-expressing Gpd2-mNeonGreen (green) and Pex14-mCherry (magenta) grown in YES or post-shift to EMM for the indicated time periods. (**B**) Maximum Z-projection spinning disk confocal images of *S. japonicus pex11*Δ cells co-expressing Gpd2-mNeonGreen (green) and Pex14- mCherry (magenta) grown in YES or post-shift to EMM for the indicated time periods. Cells outlined by a dotted line contain no discernible peroxisomes. (**A**-**B**) Scale bars represent 5 μm. Brightfield images are shown to the right of the corresponding fluorescence channels.

### Optimal lysine and histidine biosynthesis imposes an upper limit on peroxisome size

In line with the function of Pex11 in promoting peroxisome fission (Li and Gould, 2002), we observed fewer peroxisomes that were, on average, larger in *pex11*Δ *S. japonicus* cells, as compared to the wild type (Fig. 4A-C).

To investigate the role of peroxisome size further, we identified an *S. japonicus* mutant harbouring a single amino acid substitution from tryptophan to alanine at amino acid residue 224 in Pex5 (Pex5-W224A, see sequence alignment of fission yeast Pex5 proteins in Fig. S4A), which exhibited higher number of peroxisomes, coinciding with reduction in individual compartment size (Fig. 4A-C). Despite these differences in compartment organisation, the peroxisomal import of either PTS1- or PTS2-type cargoes was not affected (Fig. S4B). *pex5-W224A S. japonicus* mutant cells could grow in the minimal medium, indicating that smaller peroxisomes were capable of supporting lysine and histidine biosynthesis (growth rate of *pex5-W224A* cells in EMM was 149.2±12.3 minutes versus 162.4±15.4 minutes for wild type cells, shown as mean±SD, n=3, p-value=0.3073, unpaired t-test).

Of note, the introduction of *pex5-W224A* mutation to *pex11*Δ genetic background restored the number and the size of individual peroxisomes to that of the wild type (Fig. 4A-C). Strikingly, *pex11*Δ *pex5-W224A* double mutant cells exhibited the wild-type like growth pattern in the minimal medium (Fig. 4D), suggesting that the compartment architecture rather than Pex11 function per se defined the amino acid biosynthetic capacity of peroxisomes.

Interestingly, using 3D segmentation analysis in the wild type and the peroxin mutants described above, we observed that on a population level the peroxisome size exhibited positive correlation with an average intensity of peroxisomal proteins (see Fig. 4E for Gpd2-mNeonGreen and Fig. S4H for GFP-Lys3 and GFP-His2). This suggested that the concentration of enzymes could be considerably higher in enlarged peroxisomes in *S. japonicus pex11*Δ cells, as compared to the wild type.

An increase in enzyme abundance does not necessarily guarantee a higher biosynthetic output (Diamanti et al., 2021), especially if several pathways share a common limiting factor. We reasoned that selectively moving either Lys3 or His2 reactions out of the peroxisomal matrix to cytosol will allow us to test if peroxisome size impacted on the productivity of lysine and histidine biosynthesis. To this end, we used a GFP-binding nanobody approach, where we tethered either enzyme to the microtubule-and polar cortex-associated protein Tea1 (Behrens and Nurse, 2002) tagged with mCherry and the GFP-binding protein (GBP) (Rothbauer et al., 2006) (Fig. S4C). As a control, we constructed a truncated version of Tea1, which did not interact with microtubules or cell polarity factors and exhibited diffuse cytosolic localization (Tea1^1-407^-mCherry-GBP; Fig. S4D). We reasoned that upon the interaction with GFP-tagged Lys3 or His2, the full length Tea1-mCherry-GBP will be able to retain at least a fraction of these enzymes in the cytosol allowing them to use cytosolic NAD^+^, whereas the truncated Tea1^1-407^-mCherry-GBP will be imported into peroxisomes alongside them. The peroxisomal features, such as their size, number and enzyme intensities, were comparable in cells expressing either the full-length Tea1-mCherry GBP or the truncated Tea1^1-407^-mCherry-GBP, both in the wild type or *pex11*Δ genetic background (Fig. S4E-H and Fig. S4I-L).

The full length Tea1-mCherry-GBP colocalized with GFP-Lys3 or GFP-His2 in puncta that were often enriched at the cell tips in the minimal medium (Fig. 4F; note that a cell tip localization was more obvious in the case of Lys3). Gpd2-mTagBFP2 was found at many but not all the puncta (Fig. S4N). These results suggested that Tea1 continued to interact with the cellular polarity machinery, and GFP-Lys3 (or GFP-His2) Tea1-mCherry-GBP complexes largely decorated the surface of peroxisomes. Strikingly, the *pex11*Δ cells where either GFP-Lys3 or GFP-His2 were tethered away from the peroxisomal matrix by the full-length Tea1-mCherry-GBP, were able to sustain normal polarized growth upon the shift to the minimal medium, a rescue of the growth arrest displayed by *pex11*Δ cultures (Fig. 4G). In contrast, the introduction of the truncated Tea1^1-407^-mCherry-GBP in either GFP-Lys3 or GFP-His2 expressing *pex11*Δ cells failed to support their growth in EMM (Fig. S4M).

We concluded that peroxisome size was an important determinant in supporting the productivity of lysine and histidine biosynthesis reactions competing for the common co-factor NAD^+^ (Fig. 5).

**Figure 4.**
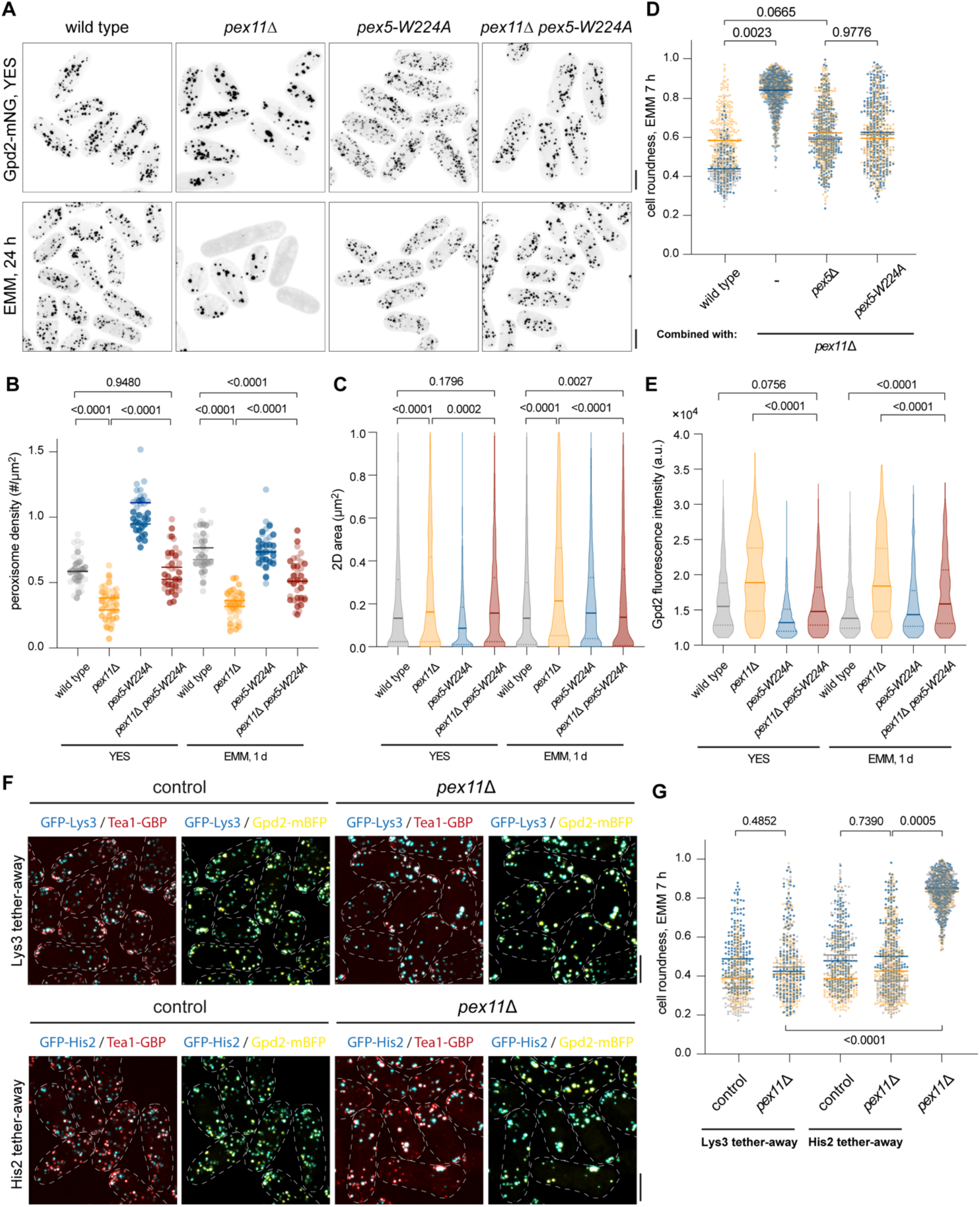
Productivity of lysine and histidine biosynthesis relies on optimal peroxisome size. (A) Maximum Z-projection spinning disk confocal images of *S. japonicus* cells of indicated genotypes expressing Gpd2-mNeonGreen grown in YES or post-shift to EMM for 24 hours. (B) Density of peroxisome compartments (number # per μm^2^) from cell populations shown in (**A**). (**C**) Size of peroxisomes (expressed as Z-projected 2D area in μm^2^) from cell populations shown in (**A**). (**D**) Cell morphology profiles (using a ‘cell roundness’ metric) of *S. japonicus* cells of indicated genotypes sampled at 7 hours post-shift to EMM. (**E**) Average fluorescence intensities of Gpd2-mNeonGreen within peroxisome compartments of cell populations shown in (**A**). (**F**) Colour overlays of maximum Z-projection spinning disk confocal images of *S. japonicus* wild type (left) and *pex11*Δ cells (right) upon the shift to EMM for 24 hours. These cells co-express Tea1-mCherry-GBP (red) and Gpd2-mTagBFP2 (yellow) in combination with either GFP-Lys3 (cyan, upper panel) or GFP-His2 (cyan, lower panel). Cell boundaries are outlined. (**G**) Cell morphology profiles (using a ‘cell roundness’ metric) of *S. japonicus* cells of indicated genotypes sampled at 7 hours post-shift to EMM. (**A, F**) Scale bars represent 5 μm. (**B**, **C**, **E**) Values are derived from two biological replicates. *p* values are derived from unpaired Welch’s t-test. Solid lines represent population medians. Dotted lines represent the upper and lower quartiles. (**D**, **G**) Values are derived from three biological replicates. *p* values are derived from unpaired t-test. Bars represent population medians.

**Supplemental Figure 4.**
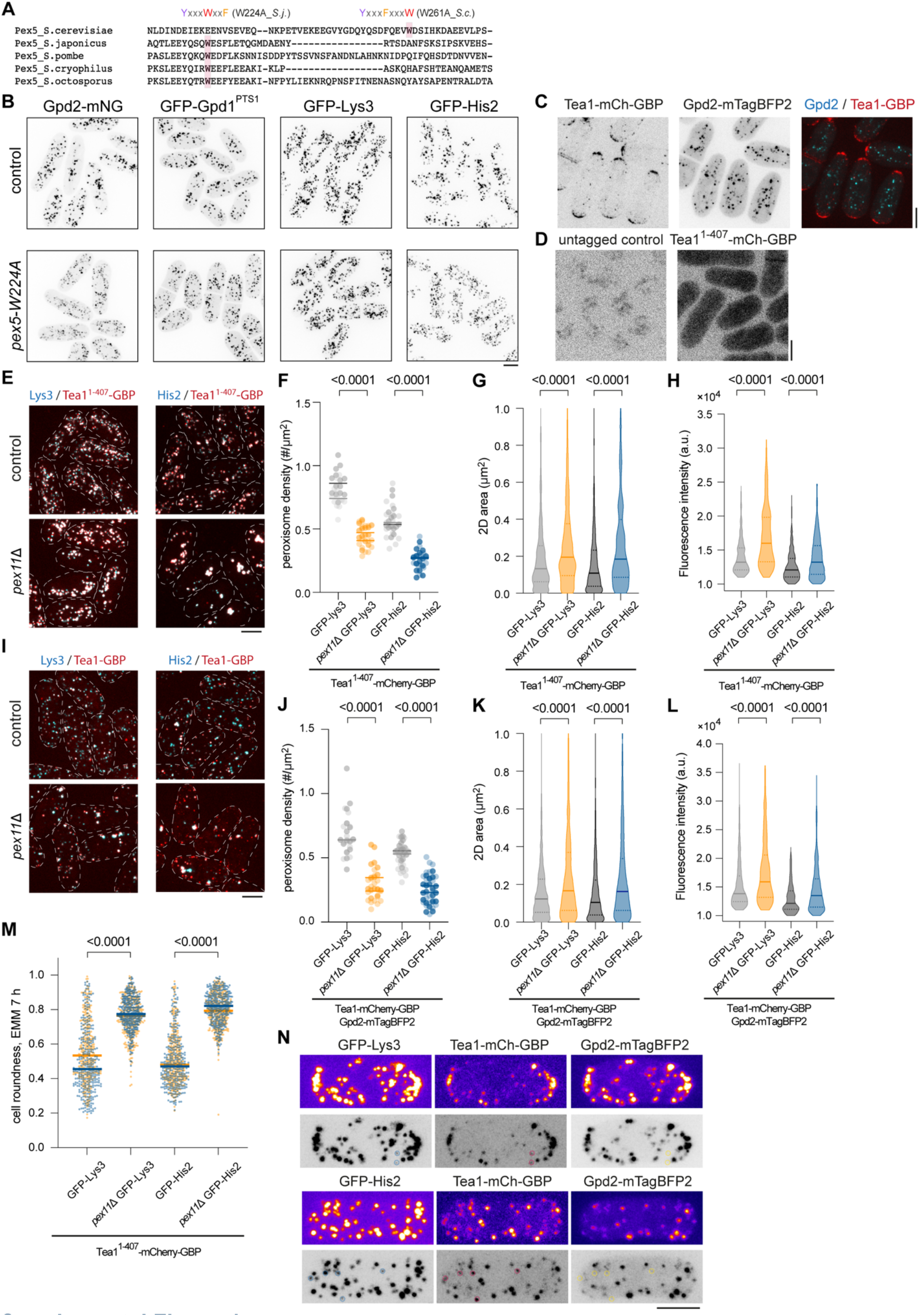
**(A)** MUSCLE alignment of peptide sequences spanning the YxxxWxxF motif conserved in Pex5 orthologs from the fission yeast clade, together with the YxxxFxxxW motif in the *S*. *cerevisiae* Pex5. (**B**) Maximum Z-projection spinning disk confocal images of wild type and *pex5*-*W224A S. japonicus* cells expressing Gpd2-mNeonGreen, GFP-Gpd1^PTS1^, GFP-Lys3 or GFP-His2. (**C**) Maximum Z-projection spinning disk confocal images of *S. japonicus* cells co-expressing Tea1-mCherry-GBP and Gpd2-mTagBFP2, shown as individual channels in grey, together with a colour overlay. (**D**) Maximum Z-projection spinning disk confocal images of untagged wild type *S. japonicus* cells (*left*) and cells expressing Tea1^1-407^-mCherry-GBP (*right*). **(A)** (**E**) Colour overlays of maximum Z-projection spinning disk confocal images of *S. japonicus* control (*top*) and *pex11*Δ cells (*bottom*) grown in YES, co-expressing Tea1^1-407^-mCherry-GBP (red) with GFP-Lys3 (cyan) or GFP-His2 (cyan). (**F**) Density of peroxisome compartments (number # per μm^2^) from cell populations shown in (**E**). (**G**) Size of peroxisomes (expressed as Z-projected 2D area in μm^2^) from cell populations shown in (**E**). (**H**) Average fluorescence intensities of either GFP-Lys3 or GFP-His2 labelled entities of cell populations shown in (**E**). **(I)** Colour overlays of maximum Z-projection spinning disk confocal images of *S. japonicus* control (*top*) and *pex11*Δ cells (*bottom*) grown in YES, co-expressing Tea1-mCherry-GBP (red) with GFP-Lys3 (cyan) or GFP-His2 (cyan). (**F**) Density of peroxisome compartments (number # per μm^2^) from cell populations shown in (**I**). (**G**) Size of peroxisomes (expressed as Z-projected 2D area in μm^2^) from cell populations shown in (**I**). (**H**) Average fluorescence intensities of either GFP-Lys3 or GFP-His2 labelled entities of cell populations shown in (**I**). **(A)** (**M**) Cell morphology profiles (using a ‘cell roundness’ metric) of *S. japonicus* cells of indicated genotypes sampled at 7 hours post-shift to EMM. (**N**) Maximum Z-projection spinning disk confocal images of *S. japonicus* cells co-expressing Tea1-mCherry-GBP and Gpd2-mTagBFP2 with either GFP-Lys3 (*top*) or GFP-His2 (*bottom*), shown as individual channel in pseudo-colour and grey. Circles highlight co-localization between GFP-tagged enzymes with Tea1-mCh-GBP without visible signal from Gpd2-mTagBFP2. (**B**-**E**, **I**, **N**) Scale bars represent 5 μm. (**F**-**H, J**-**M**) Values are derived from two biological replicates. *p* values are derived from Welch’s unpaired t-test. Solid bars represent population medians. Dotted bars represent the upper and lower quartiles.

**Figure 5.**
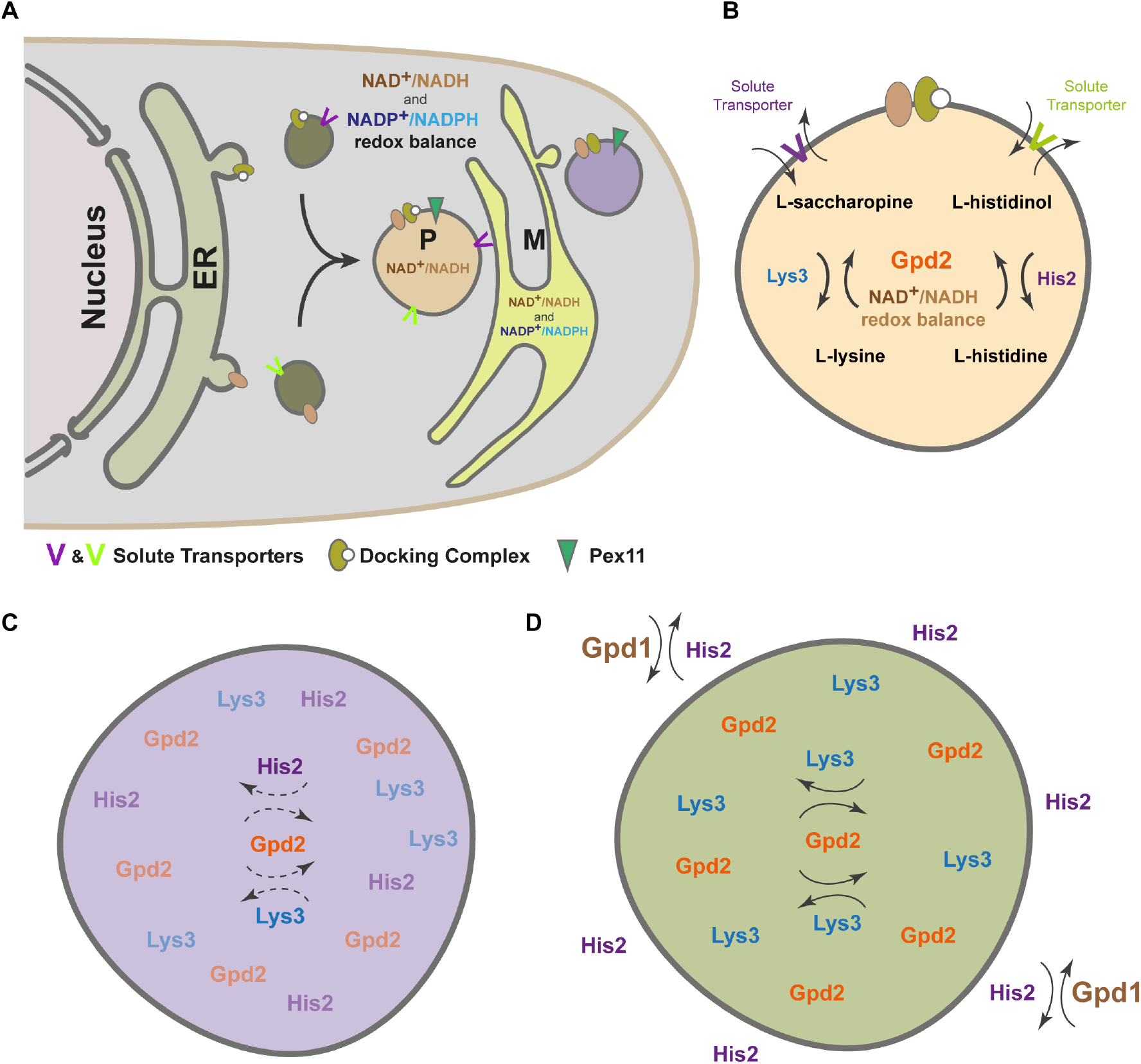
A pictorial model of intracellular redox balance facilitated by compartmentalization of amino acid biosynthesis enzymes within optimally sized peroxisomes. **(A)** Multiple NAD^+^/NADH or NADP+/NADPH-dependent metabolic reactions are present in the cytosol or compartmentalized within organelles, such as peroxisomes. Peroxisome biogenesis consists of two pathways, de novo biogenesis from the ER or the fission of pre-existing mature peroxisomes mediated by Pex11. (**B**) Glycerol-3-phosphate dehydrogenase Gpd2 produces NAD^+^ for histidine and lysine biosynthesis in peroxisomes. (**C**) Lysine and histidine production decrease in densely packed large peroxisomes in *pex11*Δ cells. (**D**) Relieving competition between compartmentalized reactions restores biosynthetic capability in enlarged peroxisomes of *pex11*Δ cells.

## Discussion

Vegetatively growing *S. japonicus* maintains a large number of peroxisomes, both in rich and minimal media. The overall peroxisome biogenesis machinery in this organism is well conserved (Jansen et al., 2021; Rutherford et al., 2022), with Pex3 being a critical upstream biogenesis factor, Pex5 and Pex7 serving as PTS1 and PTS2 import cargo receptors (Fig. 2 and Fig. S2), respectively, and Pex11 promoting peroxisomal fission (Fig. 4). Interestingly, *S. japonicus* appears to rely on Pex5 to facilitate Pex7-dependent cargo import (Fig. S2A). This is similar to many other organisms, including mammals and plants. Of note, the Schizosaccharomyces genomes do not encode the Pex7 co-receptors, used in budding yeast and Neurospora lineages (Jansen et al., 2021).

Compartmentalization of the NAD^+^ dependent reactions producing lysine and histidine appears to be a critical function for fission yeast peroxisomes. Unlike *S. cerevisiae*, the fission yeasts do not have the malate/oxaloacetate NAD^+^/NADH Mdh3 shuttle (Al-Saryi et al., 2017), and rely fully on Gpd2 to oxidize NADH in this compartment. The release of Lys3 and His2 enzymes into cytosol by genetically ablating Pex5 leads to a pronounced disbalance in cellular concentrations of lysine and histidine, but also a number of other amino acids. Of note, although histidine levels in *pex5*Δ *S. japonicus* cells drop in comparison to the wild type, the concentration of lysine is increased virtually four-fold. In *S. pombe* and *S. cerevisiae*, an increase in free lysine inhibits the first step in the lysine biosynthesis pathway through the homocitrate synthase ((Bulfer et al., 2009; Bulfer et al., 2010; Feller et al., 1999); Fig. S2D). Interestingly, the *S. japonicus* homocitrate synthase Lys4 localizes to mitochondria (Fig. S2E), unlike its homologs in *S. pombe* and *S. cerevisiae*, which are distributed throughout the cytosol and/or the nucleus (Fig. S2G and (Breker et al., 2014; Breker et al., 2013; Chen et al., 1997; Feller et al., 1999)). Compartmentalization may shield the *S. japonicus* Lys4 from such an inhibition (Alam et al., 2017); it is also possible that it had acquired mutation(s) desensitizing it to inhibition by the end product of the pathway (Isogai et al., 2021).

In the absence of negative feedback over lysine production (Feller et al., 1994; Feller et al., 1999; Ramos et al., 1988), peroxisomal compartmentalization of the lysine producer Lys3 may prevent it from competing for the common NAD^+^ pool in the cytosol. Of note, although extra GPD activity in the cytosol does partially ameliorate the disbalance in the overall free amino acid profile of *pex5*Δ *S. japonicus* cells, it boosts the lysine levels even further, emphasizing the importance of peroxisomal sequestration of Lys3 enzymatic activity (Fig. 2D, E, G).

Subcellular compartmentalization may allow better local management of redox balance risks. For instance, we notice that the levels of proline and threonine, which are synthesized in the cytosol in a NADPH-dependent manner decrease in *pex5*Δ cells, as compared to the wild type (Fig. 2D). Yet, we do not observe any differences in the levels of valine and isoleucine, which also require NADPH for their synthesis but are produced in mitochondria. This phenomenon could be related to an increased flux through the entire lysine biosynthesis pathway, since the NADPH-consuming part of it localizes in the cytosol (Fig. S2D, E).

Cellular compartmentalization has emerged across all kingdoms of life. Presumably, it provides a universal strategy to modulate flux of biological networks by spatially segregating enzymes and their coupled reactants into organelles, whether enclosed by membranes and proteinaceous shells, or phase separated into condensates (Alam et al., 2017; Pareek et al., 2021). Theoretical work has suggested several principles dictating the overall productivity of compartmentalized enzymatic reactions (Hinzpeter et al., 2017). It was suggested that the optimal enzyme density in the compartment is far below the maximal occupancy, and that each reaction compartment has a critical size, above which the productivity drops (Hinzpeter et al., 2017). Therefore, for distributed organelles such as peroxisomes, an optimal compartmentalization strategy would be establishing multiple compartments, each under a critical size, and having less than maximal enzyme packing (Hinzpeter et al., 2017). Our results showing that the abnormally large peroxisomes overpacking metabolic enzymes in *S. japonicus pex11*Δ cells are functionally deficient in compartmentalized amino acid biosynthesis (Fig. 3 and Fig. 4), support this hypothesis. It will be of interest to investigate if there exist the mechanism(s) controlling peroxisomal cargo import and fission to optimize relative enzyme concentrations and/or peroxisome size in various physiological situations.

In a densely packed macromolecular environment of peroxisomes, a number of considerations including reactant channelling between biochemically linked enzymes, diffusibility of metabolic intermediates and allosteric inhibition, may all contribute to determining an optimal compartment size. Our work begins to illuminate the fundamental rules for enzymatic compartment organization, which could be applicable to other compartmentalized biochemistry in biological and man-made systems.

## Acknowledgements

We are grateful to the Oliferenko lab and Greg Jedd for discussions and Eugene Makeyev for suggestions on the manuscript. Many thanks to James MacRae and James Ellis (Francis Crick Institute Metabolomics Science Technology Platform) for assistance with metabolomics experiments. We are grateful to Damien Coudreuse (IGDR Bordeaux) and Aleksandar Vjestica (University of Lausanne) for sharing cloning vectors. This research was funded in whole or in part, by the Wellcome Trust (103741/Z/14/Z; 220790/Z/20/Z) and BBSRC (BB/T000481/1). For the purpose of Open Access, the author has applied a CC-BY public copyright licence to any Author Accepted Manuscript version arising from this submission.

## Author contributions

Ying Gu conceived, performed and interpreted all cell biology, genetics and physiology experiments; generated strains; analysed data; and co-wrote the manuscript. Sara Alam generated and analysed metabolomics data and edited the manuscript. Snezhana Oliferenko conceived and interpreted experiments, co-wrote and edited the manuscript.

## Materials and Methods

### Fission yeast cell culture and transformation

Standard fission yeast growth media and conditions were used for culturing *S. japonicus* and *S. pombe* (Petersen and Russell, 2016). All yeast strains in this study are listed in Supplemental Table S1, and are prototrophs, unless indicated otherwise. In medium switch experiments involving growth curve measurements or image recording, cells were first grown in YES (yeast extract with L-histidine, L-leucine, adenine and uracil supplements) medium overnight to OD_595nm_ 0.4-0.6 at 30 °C. Cells were briefly centrifuged and washed with either YES or EMM twice before re-suspending in the desired medium at OD_595nm_ 0.1-0.15. Cultures were maintained at 30 °C. Cells were sampled at indicated time points for analysis. Population growth rates were represented as the maximal doubling time calculated from optical density measurements at OD_595nm_ in exponential phase cell populations. Fission yeast transformation procedure was based on an electroporation protocol described in (Aoki et al., 2010).

### Genetic manipulation of fission yeasts

Molecular genetics manipulations were performed using PCR (Bähler et al., 1998)- or plasmid– based homologous recombination (Keeney and Boeke, 1994). All plasmids were built with the restriction enzyme method, using plasmids carrying *S. japonicus ura4*, *kanR* or *hphR* cloned into the pJK210-backbone, pSO550 (Yam et al., 2011), pSO820 (Gu and Oliferenko, 2019) or pDCJ07 (gift from D. Coudreuse). mTagBFP2 was PCR-amplified from pAV0471 (gift from A. Vjestica) and cloned into pSO820 between BamHI and NotI restriction enzyme sites. To build the peroxisomal targeting Gpd1^PTS1^, nucleotide sequences of PTS1 derived from the last 14 amino acid residues of the *S. japonicus* Lys3 was fused in frame with the *gpd1* ORF at the C-terminus. To delocalize Gpd2 from peroxisomes, fusion PCR was carried out to remove nucleotide sequences encoding the first 28 amino acid residues from the N-terminus of Gpd2, giving rise the Gpd2^PTS2Δ^. To generate point mutations in the Gpd2 PTS2 (Gpd2^3KALG^), fusion PCR was carried out using mutagenic primers containing sequences that convert encoded lysine residues at locations 4, 8 and 18 to alanine, leucine residues at locations 9, 12 and 16 to glycine and isoleucine residue at position 26 to glycine. To generate plasmid harbouring point mutation in *pex5* ORF, fusion PCR was carried out to incorporate nucleotide sequences that mutate the tryptophan residue at position 224 to alanine (*pex5*-W224A). All proteins were expressed under their native regulatory elements, with the exception of Gpd1^PTS1^, which was expressed from a *gpd2* promoter as an ectopic copy integrated into the *ura4* locus. All constructs were sequenced for verification. All primers for cloning and genotyping are shown in Supplemental Table S2.

### Microscopy

Prior to imaging, 1 mL cell culture was concentrated to 50 µL by centrifugation at 1500 g, 30 sec. 2 µL cell suspension was loaded under a 22 x 22 mm glass coverslip (VWR, thickness: 1.5).

Spinning-disk confocal images were captured with Eclipse Ti-E inverted microscope fitted with Yokogawa CSU-X1 spinning disk confocal scanning unit, equipped with ILE 405nm 100mW, ILE 488nm 50mW and ILE OBIS 561nm 50mW laser lines, Nikon CFI Plan Apo Lambda 100× (N.A. = 1.45) oil objective and Andor iXon Ultra U3-888-BV monochrome EMCCD camera. Imaging was performed at 30°C with acquisition of 13 z-slices, 0.58 µm per slice, controlled by Andor Fusion software (2.3.0.36). Temperature control was achieved by the Okolab cage incubator set at 30°C.

Epifluorescence brightfield images were acquired using Zeiss Axio Observer Z1 fluorescence microscope fitted with α Plan-FLUAR 100×/1.45 NA oil objective lens (Carl Zeiss) and the Orca-Flash4.0 C11440 camera (Hamamatsu). All images were taken at the medial focal plane.

### Cell morphology analysis

Measurements of cell roundness were based on datasets consisting of brightfield images taken using Zeiss Axio Observer Z1. Images were segmented using the yeast segmentation web tool YeastSpotter (Lu et al., 2019) . Individual cell contours were obtained by selecting regions of interests (ROI) from segmented cell masks and analysed in Fiji using the shape descriptor function(Schindelin et al., 2012).

### Quantification of cellular average fluorescence intensity

Fluorescence intensity measurement of individual cells were based on datasets acquired by Andor spinning disk confocal microscope. First, all Z-slices of the brightfield channel were projected as an average intensity image. Individual cell contour was identified and separated from cell mask after segmentation of this average intensity brightfield image using YeastSpotter. Second, we generated similar projections of Z-slices of fluorescence channel. Average fluorescence intensity profiles were obtained within the cell contours. Individual values from the same cell population per biological replicate were pooled, and population means of each dataset were calculated and plotted. Heat map data is shown in Supplemental Table 3.

### Quantification of peroxisome parameters

To segment peroxisomal compartments from cells expressing various fluorescently tagged peroxisome-associated proteins, we first generated maximum intensity projections of Z-stacks of spinning disk confocal images in Fiji (Schindelin et al., 2012). Individual cell contours were manually traced with the polygon selection tool. Selected whole-cell ROIs were used to crop the regions from the corresponding maximally projected fluorescence channels. Resulting individual fluorescence images were subjected to intensity thresholding to generate mask images for peroxisomes. If peroxisomes were clustered, they were separated in masks using the watershed function. Segmented peroxisomal ROIs were applied to the Z-plane projected fluorescence images of each cell. 2D-cell area and peroxisome parameters, including size of individual peroxisomes and total counts of segmented entities were analysed in Fiji. Peroxisome density was shown as total peroxisome counts, normalized to 2D-cell area per cell.

To measure average fluorescence intensities of peroxisomal enzymes, we applied 3D segmentation in Fiji. Briefly, Z-stacks of single cell images were analysed with 3D Object Counter v2.0 plug-in. Threshold value of 11,000, minimum size filter 1 were used for Gpd2-mNeonGreen or GFP-Lys3 labelled peroxisome segmentation. Threshold value of 10,000 and minimum size filter 1 were used for GFP-His2 labelled peroxisome segmentation.

### Amino acid quantification using Gas Chromatography-Mass Spectrometry

For metabolomics experiments, cells were pre-cultured in YES overnight, washed and re-suspended in EMM and grown for 7 hours till early-exponential phase at the time of harvest. The equivalent of 2 OD_595_ was quenched by direct injection into 100% -80°C LCMS-grade methanol (Sigma Aldrich). Polar metabolite extraction was adapted from a protocol described in (Doppler et al., 2016; Vowinckel et al., 2021) Briefly, samples were extracted twice, for 15 minutes, on ice in LCMS-grade acetonitrile/methanol/water (2:2:1 v/v) with 1nmol *scyllo*-inositol internal standard per sample (Sigma Aldrich). Sample debris was then removed via centrifugation and polar metabolite extracts were dried using a SpeedVac Vacuum Concentrator. Dried extracts were phase-separated using -20°C LCMS-grade chloroform/methanol/water (1:3:3, v/v) (Sigma Aldrich).

240µl of upper, polar phase was dried in to GC-MS glass vial inserts, followed by two 30µl methanol washes. Derivatisation was performed as previously described (MacRae et al., 2013) Briefly, samples were incubated overnight in 20µl of 20mg/ml methoxyamine hydrochloride in pyridine (Sigma Aldrich). The next day, 20µl of N,O-bis(trimetylsilyl)trifluoroacetamide (BSTFA) and 1% trimethylchlorosilane (TMCS) (Sigma Aldrich) was added and samples were incubated at room temperature for at least one hour before GC-MS analysis.

Metabolites were detected using Agilent 7890B-MS7000C GC-MS in EI mode as previously described (MacRae et al., 2013). Samples were injected in a random order alongside metabolite standards and regular hexane washes. Splitless injection was performed at 270°C in a 30m + 10m x 0.25mm DB-5MS+DG column (Agilent J&W) and helium was used as the carrier gas. The oven temperature was set as follows: 70°C (two minutes), gradient of 70°C to 295°C at 12.5°C/minute, gradient of 295°C to 350°C at 25°C/minute and a three-minute hold at 350°C. Data was acquired using MassHunter version B.07.02.1938 software (Agilent Technologies).

Samples were analysed using a combination of MassHunter Workstation (Agilent Technologies) and MANIC, an updated version of the software GAVIN (Behrends et al., 2011) for metabolite identification using retention times and mass spectra and integration of target fragment ion peaks. For abundance quantification, integrals, the known amount of scyllo-inositol internal standard (1nmol) and the known abundances of a standardised metabolite mix run in parallel to samples (kindly gifted by Dr. James I. MacRae, Francis Crick Institute) were used to calculate an estimated nmol abundance of each metabolite in each sample. The formula used for calculating molar abundances as shown in Equation 1. Abundances were normalised to harvested OD_595_.

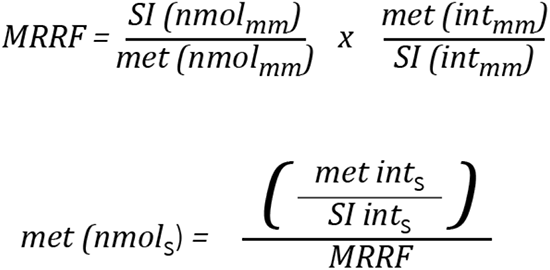

Equation 1 – Formula for nmol abundance quantification.

MRRF = molar relative response factor; SI = *scyllo*-inositol (internal standard); met = metabolite to be quantified; mm = standard metabolite mix; int = integrals; s = samples.

### NAD^+^/NADH quantification

Ratios between the oxidized form NAD^+^ and the reduced form NADH were measured using NAD^+^/NADH quantification kit (MAK037, Sigma) with whole cell lysates of wild type *S. japonicus* cells grown under indicated conditions. Briefly, early exponential cultures were harvested by centrifugation and flash freezing in liquid nitrogen. Cell pellets were then resuspended in NAD^+^/NADH extraction buffer with lysing matrix Y tubes (MP Biomedicals), containing 0.5 mm diameter zirconium oxide beads, before cells were lysed with the Fastprep-24 bead beating system (MP Biomedicals), Cell lysates were then filtered through a 10kDa protein filter (Millipore) by centrifugation at 12,000g, 4°C. Cleared extracts were processed as per the manufacturer’s instructions.

### Data visualization and statistical analysis

All data were plotted with Prism 9 (GraphPad). *p* values were calculated using indicated statistical functions in Prism 9.

## References

Agrawal, G., and Subramani, S. (2016). De novo peroxisome biogenesis: Evolving concepts and conundrums. Biochim Biophys Acta 1863, 892–901.

Al-Saryi, N.A., Al-Hejjaj, M.Y., van Roermund, C.W.T., Hulmes, G.E., Ekal, L., Payton, C., Wanders, R.J.A., and Hettema, E.H. (2017). Two NAD-linked redox shuttles maintain the peroxisomal redox balance in Saccharomyces cerevisiae. Sci Rep 7, 11868.

Alam, M.T., Olin-Sandoval, V., Stincone, A., Keller, M.A., Zelezniak, A., Luisi, B.F., and Ralser, M. (2017). The self-inhibitory nature of metabolic networks and its alleviation through compartmentalization. Nat Commun 8, 16018.

Alam, S., Gu, Y., Reichert, P., Bähler, J., and Oliferenko, S. (2022). Optimization of energy production and central carbon metabolism in a non-respiring eukaryote. bioRxiv, 2022.2012.2029.522219.

Aoki, K., Nakajima, R., Furuya, K., and Niki, H. (2010). Novel episomal vectors and a highly efficient transformation procedure for the fission yeast Schizosaccharomyces japonicus. Yeast 27, 1049–1060.

Ast, J., Bäcker, N., Bittner, E., Martorana, D., Ahmad, H., Bölker, M., and Freitag, J. (2022). Two Pex5 Proteins With Different Cargo Specificity Are Critical for Peroxisome Function in Ustilago maydis. Front Cell Dev Biol 10, 858084.

Bähler, J., Wu, J.Q., Longtine, M.S., Shah, N.G., McKenzie, A., 3rd, Steever, A.B., Wach, A., Philippsen, P., and Pringle, J.R. (1998). Heterologous modules for efficient and versatile PCR-based gene targeting in Schizosaccharomyces pombe. Yeast 14, 943–951.

Behrends, V., Tredwell, G.D., and Bundy, J.G. (2011). A software complement to AMDIS for processing GC-MS metabolomic data. Anal Biochem 415, 206–208.

Behrens, R., and Nurse, P. (2002). Roles of fission yeast tea1p in the localization of polarity factors and in organizing the microtubular cytoskeleton. J Cell Biol 157, 783–793.

Breitling, R., Sharif, O., Hartman, M.L., and Krisans, S.K. (2002). Loss of compartmentalization causes misregulation of lysine biosynthesis in peroxisome-deficient yeast cells. Eukaryot Cell 1, 978–986.

Breker, M., Gymrek, M., Moldavski, O., and Schuldiner, M. (2014). LoQAtE--Localization and Quantitation ATlas of the yeast proteomE. A new tool for multiparametric dissection of single-protein behavior in response to biological perturbations in yeast. Nucleic Acids Res 42, D726–730.

Breker, M., Gymrek, M., and Schuldiner, M. (2013). A novel single-cell screening platform reveals proteome plasticity during yeast stress responses. J Cell Biol 200, 839–850.

Bulfer, S.L., Scott, E.M., Couture, J.F., Pillus, L., and Trievel, R.C. (2009). Crystal structure and functional analysis of homocitrate synthase, an essential enzyme in lysine biosynthesis. J Biol Chem 284, 35769–35780.

Bulfer, S.L., Scott, E.M., Pillus, L., and Trievel, R.C. (2010). Structural basis for L-lysine feedback inhibition of homocitrate synthase. J Biol Chem 285, 10446–10453.

Chen, A.H., and Silver, P.A. (2012). Designing biological compartmentalization. Trends Cell Biol 22, 662–670.

Chen, S., Brockenbrough, J.S., Dove, J.E., and Aris, J.P. (1997). Homocitrate synthase is located in the nucleus in the yeast Saccharomyces cerevisiae. J Biol Chem 272, 10839–10846.

Deb, R., Ghose, S., and Nagotu, S. (2022). Increased peroxisome proliferation is associated with early yeast replicative ageing. Curr Genet 68, 207–225.

Diamanti, E., Santiago-Arcos, J., Grajales-Hernández, D., Czarnievicz, N., Comino, N., Llarena, I., Di Silvio, D., Cortajarena, A.L., and López-Gallego, F. (2021). Intraparticle Kinetics Unveil Crowding and Enzyme Distribution Effects on the Performance of Cofactor-Dependent Heterogeneous Biocatalysts. ACS Catal 11, 15051–15067.

Doppler, M., Kluger, B., Bueschl, C., Schneider, C., Krska, R., Delcambre, S., Hiller, K., Lemmens, M., and Schuhmacher, R. (2016). Stable Isotope-Assisted Evaluation of Different Extraction Solvents for Untargeted Metabolomics of Plants. Int J Mol Sci 17.

Farre, J.C., Carolino, K., Devanneaux, L., and Subramani, S. (2022). OXPHOS deficiencies affect peroxisome proliferation by downregulating genes controlled by the SNF1 signaling pathway. Elife 11.

Feller, A., Dubois, E., Ramos, F., and Pierard, A. (1994). Repression of the genes for lysine biosynthesis in Saccharomyces cerevisiae is caused by limitation of Lys14-dependent transcriptional activation. Mol Cell Biol 14, 6411–6418.

Feller, A., Ramos, F., Piérard, A., and Dubois, E. (1999). In Saccharomyces cerevisae, feedback inhibition of homocitrate synthase isoenzymes by lysine modulates the activation of LYS gene expression by Lys14p. Eur J Biochem 261, 163–170.

Galland, N., Demeure, F., Hannaert, V., Verplaetse, E., Vertommen, D., Van der Smissen, P., Courtoy, P.J., and Michels, P.A. (2007). Characterization of the role of the receptors PEX5 and PEX7 in the import of proteins into glycosomes of Trypanosoma brucei. Biochim Biophys Acta 1773, 521–535.

Gatto, G.J., Jr., Geisbrecht, B.V., Gould, S.J., and Berg, J.M. (2000). Peroxisomal targeting signal-1 recognition by the TPR domains of human PEX5. Nat Struct Biol 7, 1091–1095.

Goto-Yamada, S., Mano, S., Yamada, K., Oikawa, K., Hosokawa, Y., Hara-Nishimura, I., and Nishimura, M. (2015). Dynamics of the Light-Dependent Transition of Plant Peroxisomes. Plant Cell Physiol 56, 1264–1271.

Gu, Y., and Oliferenko, S. (2019). Cellular geometry scaling ensures robust division site positioning. Nat Commun 10, 268.

Hammond, D.J., Aman, R.A., and Wang, C.C. (1985). The role of compartmentation and glycerol kinase in the synthesis of ATP within the glycosome of Trypanosoma brucei. J Biol Chem 260, 15646–15654.

Hayashi, M., Yagi, M., Nito, K., Kamada, T., and Nishimura, M. (2005). Differential contribution of two peroxisomal protein receptors to the maintenance of peroxisomal functions in Arabidopsis. J Biol Chem 280, 14829–14835.

Hinzpeter, F., Gerland, U., and Tostevin, F. (2017). Optimal Compartmentalization Strategies for Metabolic Microcompartments. Biophys J 112, 767–779.

Isogai, S., Matsushita, T., Imanishi, H., Koonthongkaew, J., Toyokawa, Y., Nishimura, A., Yi, X., Kazlauskas, R., and Takagi, H. (2021). High-Level Production of Lysine in the Yeast Saccharomyces cerevisiae by Rational Design of Homocitrate Synthase. Appl Environ Microbiol 87, e0060021.

Jansen, R.L.M., Santana-Molina, C., van den Noort, M., Devos, D.P., and van der Klei, I.J. (2021). Comparative Genomics of Peroxisome Biogenesis Proteins: Making Sense of the PEX Proteins. Front Cell Dev Biol 9, 654163.

Jean Beltran, P.M., Cook, K.C., Hashimoto, Y., Galitzine, C., Murray, L.A., Vitek, O., and Cristea, I.M. (2018). Infection-Induced Peroxisome Biogenesis Is a Metabolic Strategy for Herpesvirus Replication. Cell Host Microbe 24, 526–541.e527.

Jourdain, I., Sontam, D., Johnson, C., Dillies, C., and Hyams, J.S. (2008). Dynamin-dependent biogenesis, cell cycle regulation and mitochondrial association of peroxisomes in fission yeast. Traffic 9, 353–365.

Keeney, J.B., and Boeke, J.D. (1994). Efficient targeted integration at leu1-32 and ura4-294 in Schizosaccharomyces pombe. Genetics 136, 849–856.

Kikuchi, Y., Kitazawa, Y., Shimatake, H., and Yamamoto, M. (1988). The primary structure of the leu1+ gene of Schizosaccharomyces pombe. Curr Genet 14, 375–379.

Kuravi, K., Nagotu, S., Krikken, A.M., Sjollema, K., Deckers, M., Erdmann, R., Veenhuis, M., and van der Klei, I.J. (2006). Dynamin-related proteins Vps1p and Dnm1p control peroxisome abundance in Saccharomyces cerevisiae. J Cell Sci 119, 3994–4001.

Legakis, J.E., Koepke, J.I., Jedeszko, C., Barlaskar, F., Terlecky, L.J., Edwards, H.J., Walton, P.A., and Terlecky, S.R. (2002). Peroxisome senescence in human fibroblasts. Mol Biol Cell 13, 4243–4255.

Li, X., and Gould, S.J. (2002). PEX11 promotes peroxisome division independently of peroxisome metabolism. J Cell Biol 156, 643–651.

Li, X., and Gould, S.J. (2003). The dynamin-like GTPase DLP1 is essential for peroxisome division and is recruited to peroxisomes in part by PEX11. J Biol Chem 278, 17012–17020.

Lu, A.X., Zarin, T., Hsu, I.S., and Moses, A.M. (2019). YeastSpotter: accurate and parameter-free web segmentation for microscopy images of yeast cells. Bioinformatics 35, 4525–4527.

MacRae, J.I., Dixon, M.W., Dearnley, M.K., Chua, H.H., Chambers, J.M., Kenny, S., Bottova, I., Tilley, L., and McConville, M.J. (2013). Mitochondrial metabolism of sexual and asexual blood stages of the malaria parasite Plasmodium falciparum. BMC Biol 11, 67.

Michels, P.A., Bringaud, F., Herman, M., and Hannaert, V. (2006). Metabolic functions of glycosomes in trypanosomatids. Biochim Biophys Acta 1763, 1463–1477.

Nakagawa, C.W., Mutoh, N., and Hayashi, Y. (1995). Transcriptional regulation of catalase gene in the fission yeast Schizosaccharomyces pombe: molecular cloning of the catalase gene and northern blot analyses of the transcript. J Biochem 118, 109–116.

Ohtsuka, H., Shimasaki, T., and Aiba, H. (2022). Response to leucine in Schizosaccharomyces pombe (fission yeast). FEMS Yeast Res 22.

Oikawa, K., Hayashi, M., Hayashi, Y., and Nishimura, M. (2019). Re-evaluation of physical interaction between plant peroxisomes and other organelles using live-cell imaging techniques. J Integr Plant Biol 61, 836–852.

Opperdoes, F.R., and Borst, P. (1977). Localization of nine glycolytic enzymes in a microbody-like organelle in Trypanosoma brucei: the glycosome. FEBS Lett 80, 360–364.

Otera, H., Okumoto, K., Tateishi, K., Ikoma, Y., Matsuda, E., Nishimura, M., Tsukamoto, T., Osumi, T., Ohashi, K., Higuchi, O., et al. (1998). Peroxisome targeting signal type 1 (PTS1) receptor is involved in import of both PTS1 and PTS2: studies with PEX5-defective CHO cell mutants. Mol Cell Biol 18, 388–399.

Pan, D., Nakatsu, T., and Kato, H. (2013). Crystal structure of peroxisomal targeting signal-2 bound to its receptor complex Pex7p-Pex21p. Nat Struct Mol Biol 20, 987–993.

Pareek, V., Sha, Z., He, J., Wingreen, N.S., and Benkovic, S.J. (2021). Metabolic channeling: predictions, deductions, and evidence. Mol Cell 81, 3775–3785.

Petersen, J., and Russell, P. (2016). Growth and the Environment of Schizosaccharomyces pombe. Cold Spring Harb Protoc 2016, pdb.top079764.

Petersen, J.G., and Holmberg, S. (1986). The ILV5 gene of Saccharomyces cerevisiae is highly expressed. Nucleic Acids Res 14, 9631–9651.

Pineda, E., Thonnus, M., Mazet, M., Mourier, A., Cahoreau, E., Kulyk, H., Dupuy, J.W., Biran, M., Masante, C., Allmann, S., et al. (2018). Glycerol supports growth of the Trypanosoma brucei bloodstream forms in the absence of glucose: Analysis of metabolic adaptations on glycerol-rich conditions. PLoS Pathog 14, e1007412.

Ramos, F., Dubois, E., and Piérard, A. (1988). Control of enzyme synthesis in the lysine biosynthetic pathway of Saccharomyces cerevisiae. Evidence for a regulatory role of gene LYS14. Eur J Biochem 171, 171–176.

Reddy, J.K., Azarnoff, D.L., and Hignite, C.E. (1980). Hypolipidaemic hepatic peroxisome proliferators form a novel class of chemical carcinogens. Nature 283, 397–398.

Reddy, J.K., and Krishnakantha, T.P. (1975). Hepatic peroxisome proliferation: induction by two novel compounds structurally unrelated to clofibrate. Science 190, 787–789.

Rhind, N., Chen, Z., Yassour, M., Thompson, D.A., Haas, B.J., Habib, N., Wapinski, I., Roy, S., Lin, M.F., Heiman, D.I., et al. (2011). Comparative functional genomics of the fission yeasts. Science 332, 930–936.

Rothbauer, U., Zolghadr, K., Tillib, S., Nowak, D., Schermelleh, L., Gahl, A., Backmann, N., Conrath, K., Muyldermans, S., Cardoso, M.C., et al. (2006). Targeting and tracing antigens in live cells with fluorescent nanobodies. Nat Methods 3, 887–889.

Rutherford, K.M., Harris, M.A., Oliferenko, S., and Wood, V. (2022). JaponicusDB: rapid deployment of a model organism database for an emerging model species. Genetics 220.

Schindelin, J., Arganda-Carreras, I., Frise, E., Kaynig, V., Longair, M., Pietzsch, T., Preibisch, S., Rueden, C., Saalfeld, S., Schmid, B., et al. (2012). Fiji: an open-source platform for biological-image analysis. Nat Methods 9, 676–682.

Veenhuis, M., Keizer, I., and Harder, W. (1979). Characterization of peroxisomes in glucose-grown Hansenula polymorpha and their development after the transfer of cells into methanol-containing media. Archives of Microbiology 120, 167–175.

Veenhuis, M., Mateblowski, M., Kunau, W.H., and Harder, W. (1987). Proliferation of microbodies in Saccharomyces cerevisiae. Yeast 3, 77–84.

Vowinckel, J., Hartl, J., Marx, H., Kerick, M., Runggatscher, K., Keller, M.A., Mülleder, M., Day, J., Weber, M., Rinnerthaler, M., et al. (2021). The metabolic growth limitations of petite cells lacking the mitochondrial genome. Nat Metab 3, 1521–1535.

Waterham, H.R., Ferdinandusse, S., and Wanders, R.J. (2016). Human disorders of peroxisome metabolism and biogenesis. Biochim Biophys Acta 1863, 922–933.

Williams, C., Opalinski, L., Landgraf, C., Costello, J., Schrader, M., Krikken, A.M., Knoops, K., Kram, A.M., Volkmer, R., and van der Klei, I.J. (2015). The membrane remodeling protein Pex11p activates the GTPase Dnm1p during peroxisomal fission. Proc Natl Acad Sci U S A 112, 6377–6382.

Woodward, A.W., and Bartel, B. (2005). The Arabidopsis peroxisomal targeting signal type 2 receptor PEX7 is necessary for peroxisome function and dependent on PEX5. Mol Biol Cell 16, 573–583.

Yam, C., He, Y., Zhang, D., Chiam, K.H., and Oliferenko, S. (2011). Divergent strategies for controlling the nuclear membrane satisfy geometric constraints during nuclear division. Curr Biol 21, 1314–1319.

Yamada, H., Ohmiya, R., Aiba, H., and Mizuno, T. (1996). Construction and characterization of a deletion mutant of gpd2 that encodes an isozyme of NADH-dependent glycerol-3-phosphate dehydrogenase in fission yeast. Biosci Biotechnol Biochem 60, 918–920.

